# Mechanism and key RNA determinants of SARS-CoV-2 Nsp1-induced endonucleolytic cleavage of mRNA

**DOI:** 10.64898/2026.06.18.733267

**Authors:** Irina S. Abaeva, Atala B. Jena, Christopher U. T. Hellen, Tatyana V. Pestova

**Affiliations:** Department of Cell Biology, SUNY Downstate Health Sciences University, Brooklyn, NY, USA

**Keywords:** SARS-CoV-2 Nsp1, eIF3g, endonucleolytic cleavage, eukaryotic translation initiation, 40S ribosomal subunit, transesterification

## Abstract

SARS-CoV-2 nonstructural protein 1 (Nsp1) binds to 40S ribosomal subunits and induces host protein synthesis shut off by inhibiting translation initiation and triggering endonucleolytic cleavage of cellular mRNAs. Irrespective of the mode of initiation, Nsp1-mediated cleavage is induced by the cooperative action of the N-terminal domain of Nsp1, the RRM domain of eIF3g and 40S subunits. Using *in vitro* reconstitution, we determined that cleavage occurs by transesterification following intramolecular nucleophilic attack of the 2’OH of the ribose on the adjacent phosphodiester bond yielding 5’OH and 2’,3‘-cyclic phosphate termini. Cleavage requires a guanosine ∼10-22 nucleotides from the 5’ end of mRNA, occurs within a narrow window upstream of this G, is most efficient between nucleotides at positions -6/-7 and -7/-8 relative to G, and shows a preference for Pu at positions -7 or -8 which provides the 2’OH for the nucleophilic attack. Zero-length UV cross-linking of Nsp1 to nucleotides at positions -1 and -2 suggests that the critical guanosine may be recognized by Nsp1. Resistance to Nsp1-mediated cleavage of SARS-CoV-2 mRNA was ensured both by the relatively long distance between its G_23_G_24_ and the 5’end and by the preceding oligoPy stretch lacking purines at positions -7 or -8 upstream of G_23_G_24_.

## INTRODUCTION

Severe acute respiratory syndrome coronavirus type 2 (SARS-CoV-2), the etiologic agent of the COVID-19 pandemic, is a betacoronavirus in the family *Coronaviridae*. It has a ∼30 kb single-stranded positive-sense RNA genome with a capped 5’-terminus, 5’ and 3’ untranslated regions (UTRs) and a 3’poly(A) tail. The genomic RNA (gRNA) is translated to yield polyproteins that are processed by viral proteases to yield non-structural proteins Nsp1 - Nsp16. It also acts as the template for synthesis of a nested set of capped and polyadenylated subgenomic (sg) RNAs (Sola et al., 2015). This process involves template switching during transcription, and all sgRNAs consequently have a ∼70 nt-long leader sequence that is identical to the 5’-terminal region of the gRNA, followed by a variable region extending to the initiation codon that is between 1 and 230 nt long in different sgRNAs (Kim et al., 2020, 2021; Wang et al., 2021).

SARS-CoV-2 and SARS-CoV Nsp1s are major pathogenesis factors. They dampen innate immune responses, and mutations in Nsp1 impair viral replication in cells with an intact IFN response (Kamitani et al., 2006; Wathelet et al., 2007; Züst et al., 2007; Lin et al., 2021; Fisher et al., 2022). Nsp1s inhibit translation by impairing initiation and by inducing the endonucleolytic cleavage of cellular mRNAs, which are subsequently degraded by Xrn1 (Kamitani et al., 2006; Huang et al., 2011; Gaglia et al., 2012; Finkel et al., 2021; Mendez et al., 2021).

SARS-CoV-2 Nsp1 is 180 a.a. long and has a structurally conserved ∼120 a.a.-long N-terminal core (Clark et al., 2021; Semper et al., 2021; Wang et al., 2023). The C-terminal region (a.a. 126-180) is unstructured, but when bound to the 40S ribosomal subunit, a.a. 154-180 form a mini-domain consisting of two α-helices that is inserted into the entrance portion of the mRNA-binding channel (Schubert et al., 2020; Thoms et al., 2020; Yuan et al., 2020). The position of this mini-domain would clash sterically with mRNA, accounting for the mechanism of inhibition of initiation by betacoronavirus Nsp1s.

*In vitro* reconstitution revealed that Nsp1-mediated mRNA cleavage occurs following ribosomal attachment and, irrespective of the mode of initiation on any given mRNA, is induced by the cooperative action of Nsp1’s N-terminal domain (NTD) and eIF3g’s RRM domain (Abaeva et al., 2023).The ribosomal location of the RRM domain at the mRNA entrance (Brito Querido et al., 2020) implies that Nsp1-induced cleavage takes place on the solvent side of the 40S subunit downstream from the mRNA entrance. Mutational analysis identified a positively charged surface on the Nsp1 NTD and a surface on the RRM domain located over the mRNA-binding channel that contain residues that are required for cleavage (Abaeva et al., 2023). Nsp1-mediated cleavage usually takes place at several sites on an mRNA and occurs sequentially starting from the 5’end rather than randomly. The initial cleavage would remove the capped 5’-terminal region from mRNA, and even without subsequent degradation of the mRNA, this would have a profound effect on host protein synthesis.

However, many aspects of Nsp1-mediated cleavage remain poorly understood. The key outstanding questions concerning the mechanism of cleavage include the individual roles of all components (i.e. the Nsp1 NTD, eIF3g’s RRM domain and the 40S subunit) and the identity of the component(s) that is (or are) directly responsible for the formation of the nuclease active center. The nucleotide specificity of this process (if any) has also not been established. Endonucleolytic cleavage of RNA occurs either by a direct, metal ion-dependent cleavage, commonly yielding 3’OH and 5’-phosphate termini, or by metal ion-independent *trans*-esterification resulting from nucleophilic attack by 2’OH of a ribose on the neighboring phosphodiester bond to generate 5’OH and 2’,3’ cyclic phosphate termini (e.g. Yang, 2011). Which of these mechanisms is used for Nsp1-induced cleavage also remains unknown.

The multi-level mode of translation inhibition by Nsp1 raises the question of how SARS-CoV-2 mRNAs evade repression to establish a productive infection (Banerjee et al., 2020; Mendez et al.,2021; Tidu et al., 2021). The 265 nt-long 5’UTR of the gRNA comprises five domains SL1-SL5 (de Moura et al., 2024; Huston et al., 2021; Miao et al., 2021; Kretsch et al., 2024). SL5 contains the viral initiation codon AUG_265_ and the proximal segment of the Nsp1 coding region. Ribosome profiling experiments identified two conserved alternative initiation codons in the SARS-CoV-2 5’UTR: CUG_59_, located between SL2 and SL3, and AUG_107_ in SL4 (Finkel et al., 2021; Kim et al., 2021; Aviner et al., 2024). The short upstream ORF (uORF) that initiates at AUG_107_ is a conserved feature of most coronaviruses and has a mild repressive effect on SARS-CoV-2 translation (Condé et al., 2022). It is under positive genetic selection in Mouse hepatitis virus (MHV), another member of the betacoronavirus genus (Wu et al., 2014; Irigoyen et al., 2016). Initiation on SARS-CoV-2 gRNA is robust and strongly dependent on the eukaryotic initiation factor (eIF) eIF4E, eIF4A and eIF4G subunits of the eIF4F cap-binding complex (Condé et al., 2022).

The 5’-terminal SL1 of the SARS-CoV-2 5’UTR confers resistance to inhibition of translation by Nsp1 (Banerjee et al., 2020; Vora et al., 2020; Tidu et al., 2021; Bujanic et al., 2022) as previously established for SARS-CoV (Huang et al., 2011). Detailed analyses revealed the primary importance of its sequence rather than structure (Bujanic et al., 2022; Slobodin et al., 2022): resistance depended on the absence of guanosines in a specific window (nt 9-22 of the SARS-CoV-2 5’UTR) and on the proximity of this element to the 5’ end of the mRNA (Slobodin et al., 2022; Berlanga et al., 2025). Substitution of guanosines between nt 4-36 in a control nonviral 5’UTR also resulted in resistance to Nsp1-mediated repression of expression (Slobodin et al., 2022), whereas inclusion of Gs in this window, particularly in cap-proximal locations, abrogated resistance to inhibition (Tidu et al., 2021; Bujanic et al., 2022; Slobodin et al., 2022; Chen et al., 2023). Cellular mRNAs that have a G-less sequence in this region, including mRNAs that contain the 5’-terminal oligopyrimidine (TOP) motif, are resistant to Nsp1-mediated translation repression (Rao et al., 2021; Galbraith et al., 2026). Although the presence/absence of guanosines in a certain window from the 5’ end has been reported not to alter mRNA stability in either rabbit reticulocyte lysate or transfected cells, and other explanations for the effect of such changes on translation were suggested (Slobodin et al., 2022), we noticed that all Nsp1-induced cleavage sites observed on different mRNAs (Abaeva et al., 2023) occurred in a specific region upstream of guanosines that were located in this window from the 5’ end, including the first and all subsequent cleavages. Moreover, substitution of these Gs in some mutants resulted in the loss of the corresponding cleavage sites.

Here, we employed the *in vitro* reconstitution approach to investigate the mechanism of Nsp1-mediated cleavage and to identify its RNA determinants.

## RESULTS

### The critical role of guanosines and the transesterification mechanism of Nsp1-mediated RNA cleavage

To investigate the dependence of Nsp1-mediated cleavage on the presence of a guanosine at a specific distance from the 5’ end of mRNA, we employed a set of mRNAs with unstructured 5’UTRs comprising eleven CAA repeats and a single G at defined positions 8 to 24 nt. from the 5’end (Figure 1A, upper panel). To induce cleavage, mRNAs were incubated with Nsp1, 40S subunits and eIFs 3/4A/4B/4G_736-1115_. Cleavage sites were then identified by primer extension. Strong cleavage was observed on mRNAs containing Gs at positions 14 and 20 (G14 and G20) (Figure 1A, lower panel). Cleavage of mRNAs with G12, G18 and G24 was less efficient, and cleavage of mRNAs with G10, G16 and G22 was extremely weak (Figure 1A, lower panel). Efficient cleavage on mRNAs with G14 and G20 occurred between nucleotides at positions -6 and -7 relative to G, whereas less efficient cleavage of mRNAs with G12, G18 and G24 occurred between positions -7 and -8 (indicated on Figure 1A, upper panel). mRNAs that were cleaved with comparable efficiency had identical nucleotide sequences upstream and downstream of G, whereas these sequences differed between the three groups of mRNAs. Strikingly, all cleavages occurred between the PuPy pair AC, and the difference in the distance between the sites of cleavage and Gs for mRNAs with G14/G20 and mRNAs with G12/G18/G24 could therefore result from the difference in the positions of the AC pairs. In both mRNA groups, a greater separation of G from the 5’-end reduced the efficiency of cleavage. In poorly cleaved G10/G16/G22 mRNAs, the AC pairs are formed by -5/-6 and -8/-9 nucleotides, and it is plausible to suggest that AC pairs at such positions relative to G cannot be effectively accommodated into the active center of the nuclease.

**Figure 1.**
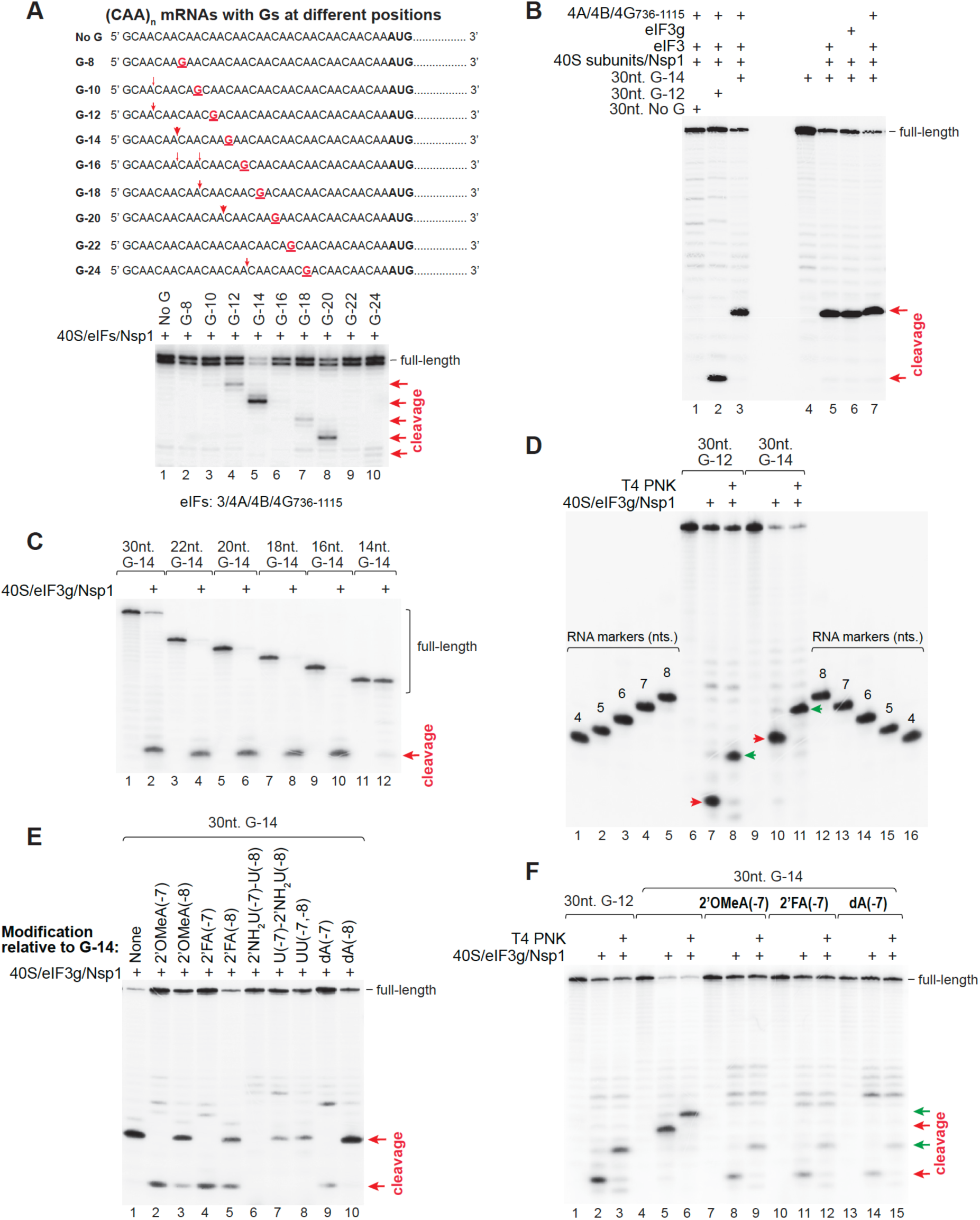
Mechanism of Nsp1-induced RNA cleavage. (A) A critical requirement for G at a specific distance from the 5’ end of mRNA for Nsp1-mediated cleavage. Upper panel - schematic representation of mRNAs containing unstructured 34nt.-long 5’UTRs comprising CAA repeats interrupted by a single G (red) at different positions from the 5’ end. Lower panel – Nsp1-mediated cleavage of mRNAs shown in the upper panel in the presence of the indicated translation components, assayed by primer extension. Cleavage sites are shown on the right and marked on 5’ UTR sequences in the upper panel (red arrows). (B) Factor requirements of Nsp1-mediated cleavage of 30nt.-long unstructured RNAs corresponding to the 5’-terminal regions of mRNAs shown in A (upper panel). After incubation of 5’[^32^P]-labeled RNAs with Nsp1 and the indicated translation components, cleavage products were visualized directly by electrophoresis (red arrows). (C) The length of RNA downstream of G required for Nsp1-induced cleavage. 14-30nt.-long 5’[^32^P]-labeled (CAA)n RNAs containing G14 were incubated with Nsp1, eIF3g and 40S subunits after which cleavage products were visualized by electrophoresis (red arrow). (D) Determination of the cleavage site and the location of the phosphate group after cleavage. 5’[^32^P]-labeled 30nt.-long (CAA)n RNAs containing G12 or G14 were cleaved by incubation with Nsp1, eIF3g and 40S subunits. Cleavage products were either treated with T4 PNK or left untreated and visualized by electrophoresis (green and red arrows, respectively). To estimate the size of the cleavage products, 4-8nt.-long 5’[^32^P]-labeled oligo RNA markers containing 3’OH were employed. (E) Nsp1-induced cleavage of 5’[^32^P]-labeled 30nt.-long (CAA)n RNAs with G14 containing the indicated 2’-derivatized nucleotides at positions -7 or -8 relative to G14. Cleavage products were visualized by electrophoresis (red arrows). (F) Comparison of Nsp1-induced cleavage products of 30nt.-long (CAA)n RNAs with G12 or G14 and of RNAs containing the indicated 2’-derivatized nucleotides at position -7 relative to G14. Cleavage products of the corresponding 5’[^32^P]-labeled RNAs were either treated with T4 PNK or left untreated and visualized by electrophoresis (green and red arrows, respectively).

To investigate the mechanism of Nsp1-mediated cleavage further, an assay system was required that could use much shorter RNAs. We therefore investigated cleavage of 30nt-long RNAs with sequences identical to nt 1-30 of the 5’UTRs of “No G”, G12 and G14 mRNAs shown in Figure 1A. RNAs were 5’[^32^P]-labeled and incubated with Nsp1, 40S subunits and various sets of initiation factors. Cleavage products were resolved directly on 20% sequencing gel and visualized by phosphor imaging. Both 30nt-long G12 and G14 RNAs were cleaved efficiently in the presence of eIFs 3/4A/4B/4G_736-1115_ (Figure 1B, lanes 2-3). Importantly, eIF3g could replace eIFs 3/4A/4B/4G_736-1115_ (Figure 1B, lane 6), which simplified the system for further biochemical studies. 3’-terminal truncations of 30nt G14 RNA revealed that efficient Nsp1-mediated cleavage required only 2 nucleotides downstream from G14 (Figure 1C).

To determine whether Nsp1 mediates RNA hydrolysis by direct cleavage yielding 3’OH and 5’phosphate termini or by transesterification, which involves nucleophilic attack of the 2’OH of the ribose on the neighboring phosphodiester bond yielding 5’OH and 2’,3‘-cyclic phosphate-linked cleavage products (e.g. Yang, 2011), we determined the location of the phosphate group after cleavage. For this, we exploited the 3’-phosphatase activity of T4 polynucleotide kinase (PNK) (Schurer et al., 2002). 5’[^32^P]-labeled 30nt-long G12 and G14 RNAs were incubated with Nsp1, 40S subunits and eIF3g and either treated with T4 PNK or left untreated. Cleavage products were then resolved on 20% sequencing gel with 4-8nt-long 5’[^32^P]-labeled RNA markers that had a 3’OH hydroxyl group and nucleotide sequences identical to the 5’-terminal sequences of G12 and G14 RNAs. Treatment with T4 PNK shifted the mobility of 5’-terminal cleavage products to a higher position due to removal of a negatively charged phosphate group (Figure 1D, lanes 7-8 and 10-11), indicating that Nsp1-mediated cleavage yields a 2’,3‘-cyclic phosphate and occurs by transesterification. The mobilities of T4 PNK-treated 5’-terminal cleavage products of G12 and G14 RNAs were identical to those of the 4nt- and 7nt-long RNA markers, which correlated with the sites of cleavage of G12 and G14 mRNAs determined by toe-printing (Figure 1A).

To confirm the crucial role of the 2’OH group of ribose in Nsp1-mediated cleavage, we investigated cleavage of 30nt.-long G14 RNAs containing 2’-derivatized nucleotides at positions -7 or -8 relative to G14. Regarding the cleavage site on G14 RNA, the nucleotide at position -7 would provide the 2’OH for the nucleophilic attack, whereas the nucleotide at -8 would be adjacent to it at the 5’ side. Substitution of A at position -7 by 2’-deoxy-adenosine (dA), 2’-O-methyl-adenosine (2’OMe-A) or 2’-Fluoro-adenosine (2’F-A) abrogated cleavage between -6/-7 nucleotides and resulted in the appearance of a lower intensity shorter 5’-terminal cleavage product (Figure 1E, lanes 2, 4 and 9). To determine the effect of the 2’-amino (2’NH_2_) modification, we had to use RNAs with 2’NH_2_-U at positions -7 and -8 because 2’NH_2_-A was unavailable. Consistent with the effect of other modifications, 2’NH_2_-U at position -7 was also not cleaved between -6/-7 nucleotides (Figure 1E, lane 6). In contrast, derivatization of the 2’ position in the nucleotide at position -8 had a much smaller effect. Like unmodified G14 RNA, such RNAs were all cleaved between -6/-7 nucleotides even though RNAs with 2’OMe-A and 2’F-A modifications additionally yielded shorter 5’-terminal products with mobilities identical to that of the products observed for RNAs with derivatized nucleotides at position -7 (Figure 1E, lanes 3 and 5). The mobility of the shorter products observed for G14 RNAs with 2’-derivatized nucleotides at position -7 with or without PNK treatment was the same as of the cleavage product for G12 RNA (Figure 1F), indicating that this weak cleavage occurred between AC nucleotides at positions -9/-10 upstream from G14.

Taken together, these data revealed that Nsp1-mediated cleavage occurs by intramolecular nucleophilic attack of the 2’OH of the ribose on the adjacent phosphodiester bond. Cleavage depends on 40S subunits and eIF3g, requires the presence of a guanosine in the RNA, occurs in a specific window upstream of that G with cleavage between -6/-7 and -7/-8 nucleotides being most efficient, and shows nucleotide preference.

### Contacts of RNA nucleotides located at the cleavage site and the critical G nucleotide with Nsp1, eIF3g and 40S subunit during Nsp1-mediated cleavage

The key open question about the mechanism of Nsp1-mediated cleavage concerns the individual roles of all essential components (i.e. Nsp1, eIF3g and the 40S subunit), which includes determination of their direct interaction with RNA. To investigate this, we identified specific RNA contacts using UV crosslinking of thio-derivatized nucleotides introduced at unique positions from the cleavage site to the critical G nucleotide in 19nt.-long G14 RNAs. Replacement of G14 by 6thio-G rendered RNA resistant to Nsp1-mediated cleavage (Figure S1A), underscoring the importance of this nucleotide but also making it impossible to determine its direct contacts by UV crosslinking. In contrast, introduction of 4thio-U at positions from -9 to +2 relative to G14 (Figure S1B) did not inhibit Nsp1-mediated cleavage (Figure S1C), indicating that such derivatives could be used for crosslinking. To stabilize binding of thioU-containing RNAs to the Nsp1/eIF3g/40S subunit complex during cross-linking, we additionally introduced dA at position -7 (Figures 2A and S1B), which almost abrogated cleavage (Figure S1C).

**Figure 2.**
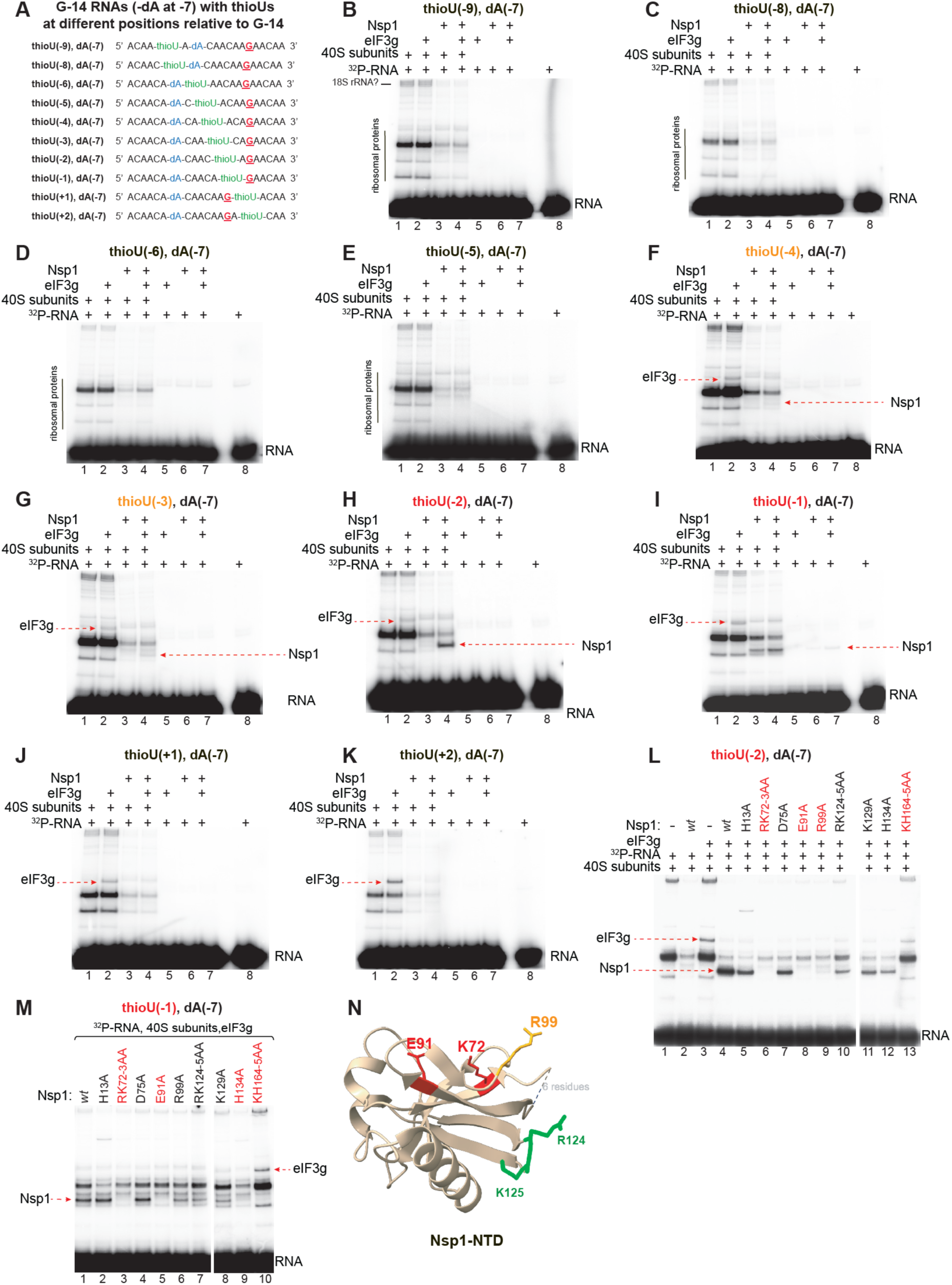
Specific protein contacts of nucleotides at different positions relative to the critical G nucleotide during Nsp1-mediated RNA cleavage. (A) Sequences of 19nt.-long (CAA)n RNAs containing G14, dA at position -7 and 4thioU at indicated positions relative to G14. (B-K) UV cross-linking of 5’[^32^P]-labeled 19nt.-long (CAA)n RNAs containing G14, dA at position -7 and 4thioU at indicated positions (panel A) with Nsp1, eIF3g and 40S subunits, assayed by SDS-PAGE and phosphor imaging. The positions of RNA and cross-linked Nsp1, eIF3g and ribosomal proteins are indicated. (L-M) UV cross-linking of 5’[^32^P]-labeled 19nt.-long (CAA)n RNAs containing G14, dA at position -7 and 4thioU at positions -2 (L) or -1 (M) with *wt* and mutant Nsp1 in the presence of 40S subunits and eIF3g, assayed by SDS-PAGE and phosphor imaging. The positions of RNA and cross-linked Nsp1 and eIF3g are indicated. (N) Ribbon diagram of the Nsp1 N-terminal domain (PDB: 7K3N) to show mutated residues in stick colored depending on their importance for cross-linking of Nsp1 to positions -1 and -2 relative to the G nucleotide: red (crucial), orange (very strong) and green (moderate) sticks.

5’[^32^P]-labeled G14 RNAs containing 4thio-U at different positions and dA at position -7 were incubated with Nsp1, eIF3g and 40S subunits individually and in different combinations and cross-linked by low-energy (360 nM) irradiation, yielding “zero-length” cross-links that represent direct contacts of thio-U nucleotides. Cross-linked products were assayed by SDS-PAGE and phosphor imaging. In the presence of 40S subunits alone, all RNAs cross-linked to identical ribosomal proteins (Figures 2B-K, lanes 1) likely due to the ability of these unstructured RNAs to associate with the mRNA-binding channel. Inclusion of eIF3g did not affect RNA cross-linking with the 40S subunits, but additional crosslinking of slightly different intensity to eIF3g was observed for RNAs containing thioU at positions from -4 to +2 relative to G14 (Figures 2F-K, lanes 2). Cross-linking of eIF3g to specific positions in RNA could reflect the reported sequence specificity of RNA binding by eIF3g (Kato et al., 2026). Nsp1 strongly reduced cross-linking of all RNAs to ribosomal proteins independently of the presence of eIF3g (Figures 2B-K, lanes 3 and 4) which is consistent with binding of Nsp1 to the mRNA-binding channel and blocking its ability to associate with RNA. It also inhibited cross-linking of RNAs to eIF3g (Figures 2F-K, lanes 4). Strikingly, Nsp1 itself was efficiently and specifically crosslinked to thio-Us at positions -1 and -2 relative to G14, and this crosslinking depended on the presence of 40S subunits and was stimulated by eIF3g to different extents (Figures 2H-I, lanes 3-4). Thus, whereas efficient cross-linking of Nsp1 to position -1 was observed even without eIF3g, cross-linking to position -2 was fully eIF3g-dependent. eIF3g-dependent low-intensity crosslinking to Nsp1 was also observed for thioUs at positions -3 and -4, with cross-linking at position -3 being slightly stronger (Figures 2F-G, lanes 3-4). No specific crosslinking was observed for thioUs surrounding the actual cleavage site (positions from -9 to -5) (Figures 2B-E).

We previously identified residues in the Nsp1-NTD and in the Nsp1 linker region that are critical for its endonuclease activity (Abaeva et al., 2023). We therefore analyzed crosslinking of corresponding Nsp1 mutants to thioU at positions -1 and -2 (Figures 2L-M). Two mutants (KR72-73AA and E91A) completely lost their ability to cross-link to thioU at both positions (Figure 2L, lanes 6 and 8; Figure 2M, lanes 3 and 5). The R99A substitution also very strongly reduced crosslinking of Nsp1 to position -2 but had a moderate effect on crosslinking to position -1 (compare Figure 2L, lane 9 with Figure 2M, lane 6). In turn, the H134A substitution nearly abolished crosslinking to position -1 but had a moderate effect on crosslinking to position -2 (compare Figure 2M, lane 9 with Figure 2L, lane 12). For both positions of thioU, the lowest effect was observed for H13A and D75A substitutions (Figure 2L, lanes 5 and 7; Figure 2M, lanes 2 and 4), whereas RK124-125AA and K129A substitutions moderately reduced crosslinking (Figure 2L, lanes 10-11; Figure 2M, lanes 7-8). Like *wt* Nsp1, these mutants all strongly reduced cross-linking of thioU-containing RNA to ribosomal proteins compared to KH164-165AA mutant Nsp1 (Figure 2L, compare lanes 4-12 with lane 13: Figure 2M, compare lanes 1-9 with lane 10) which lost the ability to bind to 40S subunits (Thoms et al., 2020; Tardivat et al., 2023). The reduced crosslinking to positions -1 and -2 was therefore not due to their inability to bind to 40S subunits. In conclusion, we found that Nsp1 makes specific direct contacts with RNA nucleotides 5’-adjacent to the critical G residue and that this RNA/Nsp1 interaction is strongly stimulated by eIF3g.

Although all tested mutants were equally inactive in RNA cleavage except for D75A mutant that possessed a low-level activity (Abaeva et al., 2023), their ability to crosslink to two RNA nucleotides adjacent to the critical G residue differed. It is plausible that KR72-73, E91 as well as R99 (substitution of which had the strongest effect on crosslinking) could be involved in recognition of the critical G nucleotide and the immediately preceding nucleotides, whereas other important residues (e.g. RK124-125, K129 or H13) could be involved in the interaction with the more upstream RNA region, in formation of the active center or in interaction with eIF3g. Consistently, in the Nsp1-NTD structure (Clarke et al., 2021; Semper et al., 2021), K72, E91 and R99 are situated in close proximity, whereas RK124-125 are further away (Figure 2N).

### The optimal nucleotide context of the cleavage site for Nsp1-mediated RNA cleavage

The data presented above suggested that Nsp1-mediated RNA cleavage possesses cleavage site nucleotide specificity. Thus, cleavage of G14 and G12 RNAs was between -6/-7 and -7/-8 nucleotides, respectively, but in both cases occurred between AC pairs (Figure 1A). To confirm that the cleavage sites were indeed determined by the positions of the AC pairs, we moved ACs one nucleotide closer to G12 and 1 nucleotide further from G14 (G12 (AC at +1) and G14 (AC at -1) mRNAs; Figure 3A, upper panel). To induce cleavage, RNAs were incubated with Nsp1, 40S subunits and eIFs 3/4A/4B/4G_736-1115_. Cleavage sites were then mapped by primer extension. As expected, changes in the positions of AC pairs altered in the positions of cleavage: G12 RNA was now cleaved between -6/-7 nucleotides, whereas G14 RNA was cleaved between -7/-8 nucleotides (Figure 3A, lower panel, compare lane 2 with lane 4 and lane 6 with lane 8). Cleavage between -6/-7 nucleotides was again more efficient than between -7/-8 nucleotides, resulting in a lower proportion of intact RNA. Substitution of the region upstream but not downstream of G in G16 RNA by corresponding regions that occur in G14 RNA (Figure 3A, upper panel) resulted in strong cleavage at position -6/-7 from G16 (Figure 3A, lower panel, lanes 9-16), indicating that very inefficient cleavage of the original G16 RNA was caused by the suboptimal positions of the upstream AC pairs rather than by the sequence located downstream from G16.

**Figure 3.**
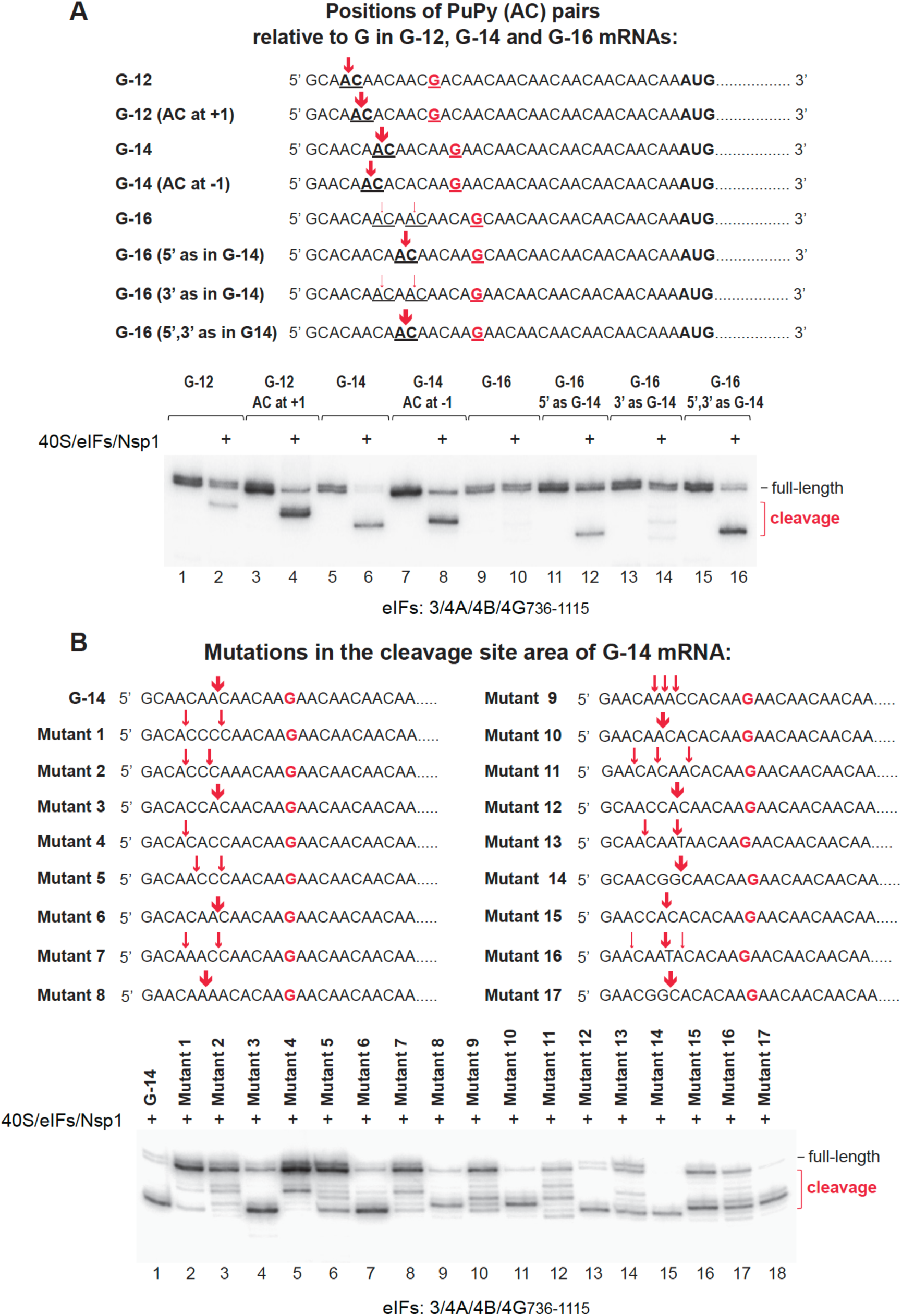
The influence of the nucleotide context of the cleavage site on the efficiency of Nsp1-mediated RNA cleavage. (A-B) Upper panels - schematic representation of 34nt.-long 5’UTRs comprising CAA repeats interrupted by G (red) at indicated positions and containing nucleotide variations at the cleavage site. Lower panels - Nsp1-mediated cleavage of mRNAs with UTRs shown on the upper panels in the presence of the indicated translation components, assayed by primer extension. Cleavage sites are shown on the right and marked on the 5’ UTRs in the upper panels (red arrows).

To investigate the nucleotide preference for Nsp1-mediated cleavage, we employed a panel of 17 RNAs containing various nucleotide sequences in the cleavage area upstream of G14 (Figure 3B, upper panel). To induce cleavage, RNAs were incubated with Nsp1, 40S subunits and eIFs 3/4A/4B/4G_736-1115_, after which cleavage sites were mapped by primer extension (Figure 3B, lower panel). The most efficient cleavage occurred between Pu and Py in PuPy pairs (e.g. AC, AU, GC) that were located at the optimal positions -6/-7 or -7/-8 relative to G14. When the positions of PuPy pairs were not optimal, weaker cleavage occurred at multiple sites either at the optimal distance (-6/-7 or -7/-8 from G) between other nucleotides, or between PuPy dinucleotides located at a suboptimal distance. These suboptimal distances were mostly further upstream from G14, and only a few weak cuts were observed at position -5/-6. Thus, the nuclease active site shows a general preference for Pu at positions -7 or -8 whose 2’OH hydroxyl group would be positioned for a nucleophilic attack on the adjacent phosphodiester bond.

### Inhibition of Nsp1-mediated cleavage by oligoPy sequences surrounding the guanosine

The reported resistance of TOP mRNAs to Nsp1-mediated translational repression (Rao et al., 2021) prompted us to investigate the influence on cleavage of the presence of oligoPy regions immediately before or after the critical G nucleotide. For oligoPy tracts preceding G, we employed a panel of mRNAs containing G14 after 12 5’-terminal pyrimidines that were either uninterrupted or interrupted by a single A nucleotide at different relative positions (Figure 4A, upper panel). To induce cleavage, mRNAs were incubated with Nsp1, 40S subunits and eIFs, and cleavage sites were then identified by primer extension. No cleavage was observed even on mRNAs that contained A at the optimal positions -7 or -8 relative to G14 (Figure 4A, lower panel). For oligoPy tracts following the critical G, different numbers of pyrimidines were introduced immediately after G (Figure 4B, upper panel). Increasing the number of pyrimidines from 1 to 6 progressively reduced the efficiency of cleavage (Figure 4B, lower panel).

**Figure 4.**
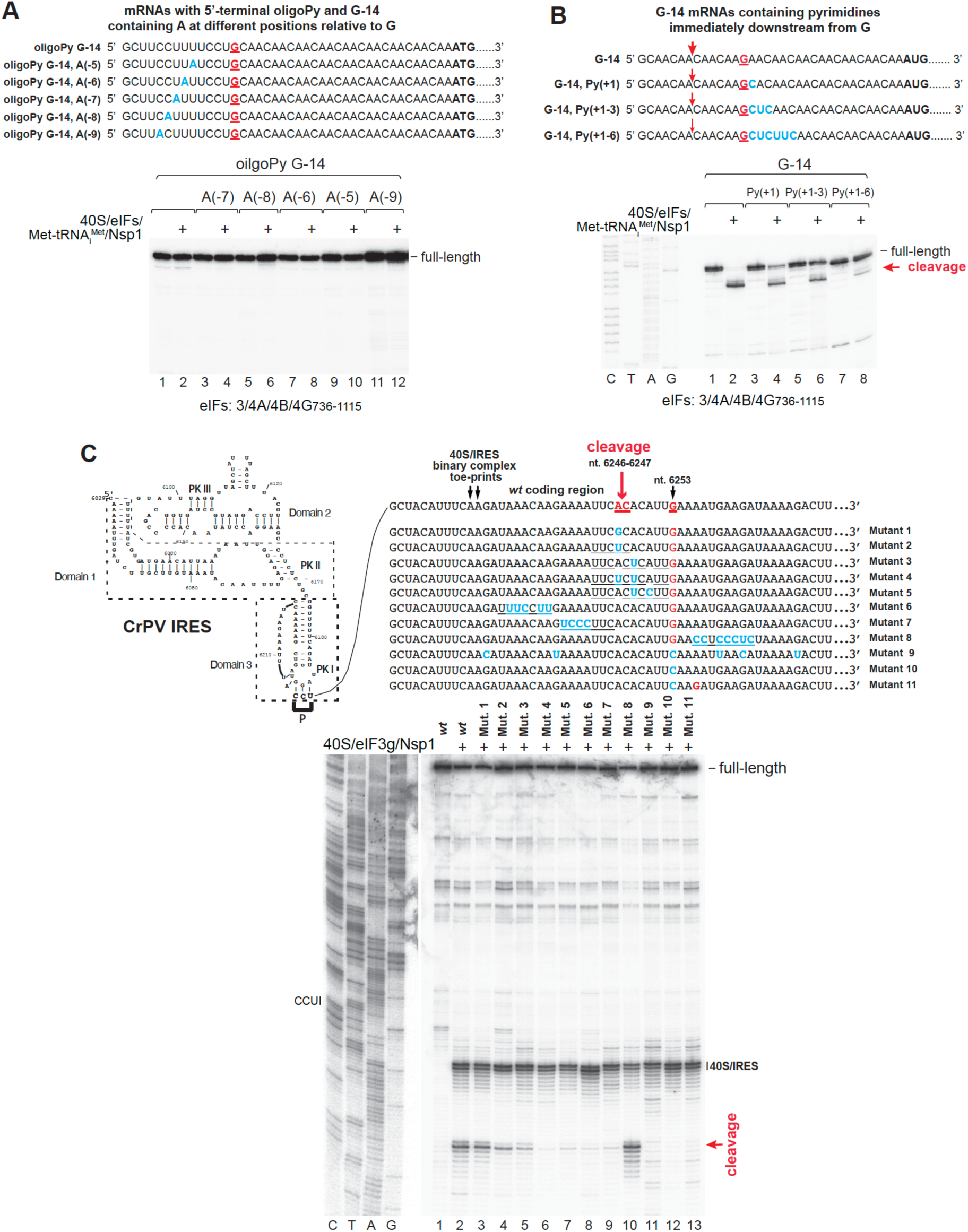
The efficiency of Nsp1-mediated cleavage of RNAs containing oligoPy regions immediately before or after the critical G nucleotide. (A-B) Upper panels – sequences of 34nt.-long 5’UTRs comprising CAA repeats interrupted by G14 (red) and containing oligoPy regions immediately before (A) or after (B) the G nucleotide. Lower panels - Nsp1-mediated cleavage of mRNAs with 5’ UTRs shown in the upper panels in the presence of the indicated translation components, assayed by primer extension. Cleavage sites are shown on the right and marked on the 5’ UTRs in the upper panels (red arrows). (C) Upper panel - Schematic representation of the CrPV IRES with the proximal cognate coding region, annotated to show IRES domains 1-3, pseudoknots PKI, PKII and PKIII, the P-site codon, the positions of toeprints of IRES-bound 40S subunits (black arrows), the critical G nucleotide (red), and the site of Nsp1-mediated cleavage (red arrow). Coding regions of mutants with introduced substitutions (blue) are shown directly below the proximal cognate coding region. Lower panel – Nsp1-mediated cleavage of *wt* and mutant CrPV IRES mRNAs shown in the upper panel in the presence of 40S subunits and eIF3g, assayed by primer extension. Toe-prints corresponding to 40S/IRES binary complexes and the cleavage site are indicated on the right. (B-C) Lanes C, T, A, and G show the corresponding sequence derived using the same primer as for primer extension. Separation of lanes by a white line in panel C indicates that these parts were juxtaposed from the same gel after different exposures.

Next, we investigated the influence of oligoPy sequences surrounding the Nsp1-mediated cleavage site on the efficiency of cleavage of mRNA containing the cricket paralysis virus (CrPV) IRES (Figure 4C, upper panel) which translates by a mechanism that differs from 5’ end-dependent scanning. This IRES binds directly to 40S subunits (Wilson et al., 2000; Pestova et al., 2004) with PKI mimicking the tRNA/mRNA interaction in the decoding center of the ribosomal A site (e.g. Schüler et al., 2006). Nsp1-mediated cleavage of CrPV IRES mRNA requires only 40S subunits and eIF3g and occurs in the coding region 18 nt downstream from the mRNA entrance between AC_6246-6247_ located at positions -6/-7 from G_6253_ (Figure 4C, upper panel; Abaeva et al., 2023). To investigate the influence of the nature of nucleotides surrounding the cleavage site, we employed a panel of mRNAs with increased Py content in this area (Figure 4C, upper panel). To induce cleavage, mRNAs were incubated with Nsp1, 40S subunits and eIF3g; cleavage sites were identified by primer extension. The G_6253_C substitution (Mutant 10) abrogated cleavage at AC_6246-6247_ as expected but did not lead to cleavage at other upstream or downstream guanosines (Figure 4C, lower panel, lane 12). Subsequent introduction of G_6256_ only 3 nt downstream from G_6253_ (Mutant 11) also did not result in Nsp1-mediated cleavage even though there was a PuPy (AC) pair at positions -7/-8 from G_6256_ (Figure 4C, lower panel, lane 13). Thus, the exact position of G in the CrPV IRES coding region relative to the mRNA entrance is extremely important for Nsp1-cleavage. As expected, substitution of A_6246_ by G (Mutant 1) did not influence Nsp1-mediated cleavage whereas the A_6246_U substitution (Mutant 2) reduced its efficiency (Figure 4C, lower panel, lanes 3-4). Enrichment of the area immediately upstream of G_6253_ and surrounding the actual cleavage site with Py residues (Mutants 3-5) further reduced the level of cleavage (Figure 4C, lower panel, lanes 5-7). Surprisingly, introduction of long oligoPy stretches even further upstream (Mutants 6 and 7) also strongly inhibited cleavage (Figure 4C, lower panel, lanes 8-9), whereas introduction of oligoPy stretch 3 nucleotides downstream from G_6253_ (Mutant 8) had no effect (Figure 4C, lower panel, lane 10).

Taken together, these data indicate that oligoPy stretches immediately upstream of the critical G and surrounding the actual cleavage site impair Nsp1-mediated cleavage irrespective of the mode of translation initiation.

### The mechanism and factor requirements for initiation on genomic and subgenomic SARS-CoV-2 mRNAs

To determine the mechanism by which SARS-CoV-2 genomic and subgenomic mRNAs outcompete cellular mRNAs in conditions of viral expression of Nsp1, we first investigated the factor-requirements and the mechanism of initiation on viral RNAs. The 5’UTRs of SARS-CoV-2 genomic (Figure 5A) and sg RNAs share a ∼70 nt-long 5’-terminal element. It forms 3 stem-loops (SL1 - SL3) and contains a near-cognate initiation codon CUG_59_ that is a site of ribosomal pausing (Finkel et al., 2021; Kim et al., 2021; Aviner et al., 2022). The initiation codon in sg RNAs is closely adjacent to the 70nt-long common element, but in genomic mRNAs, the initiation codon AUG_266_ is ∼200nt downstream. The longer genomic 5’UTR contains additional stem-loops (SL4-SL5) and an internal AUG_107_ codon linked to a short ORF which is under positive genetic selection in MHV (Wu et al., 2014) and overlaps with the short ORF that initiates at CUG_59_.

**Figure 5.**
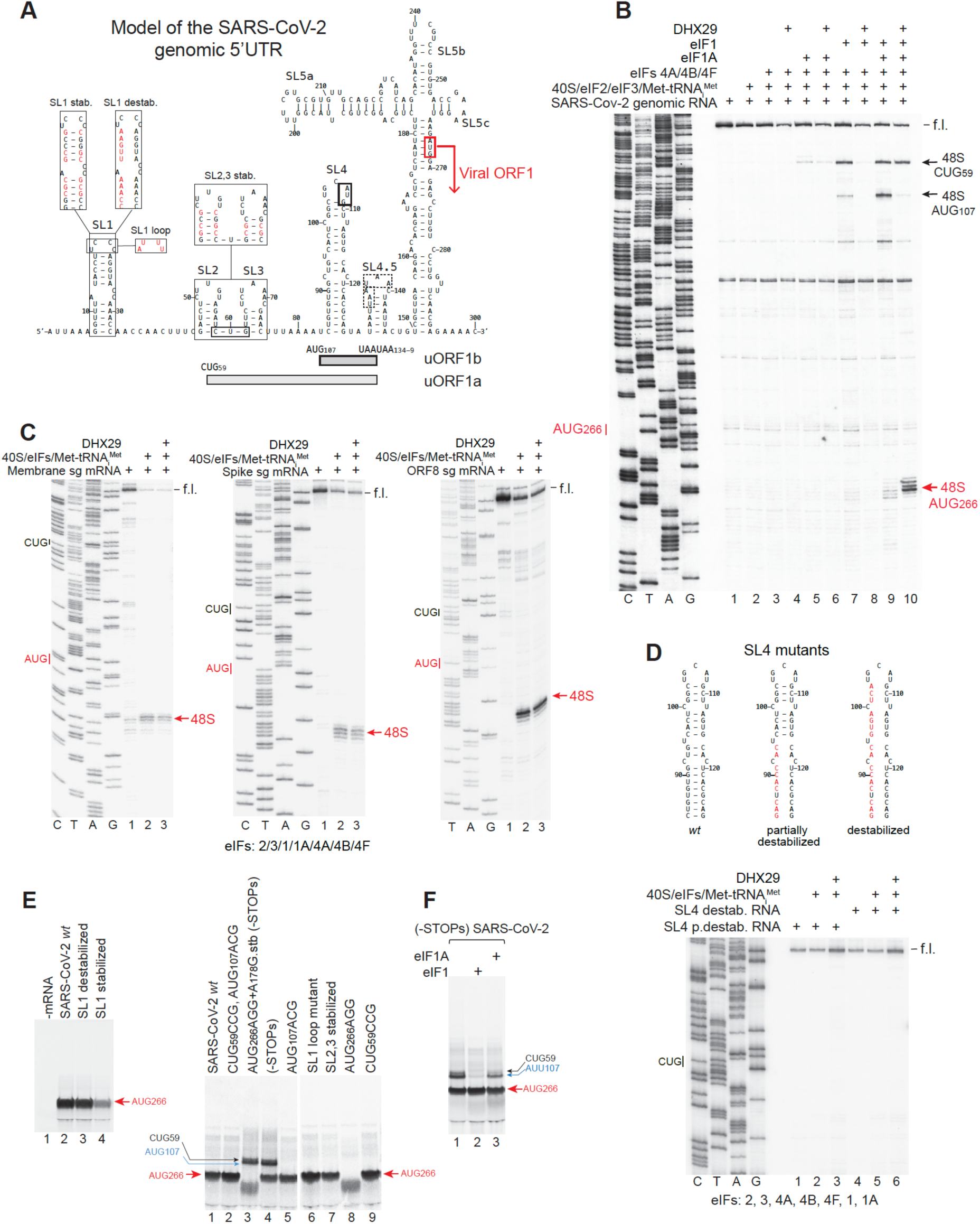
The mechanism and factor requirements for initiation on genomic and subgenomic SARS-Cov-2 mRNAs. (A) Secondary structure model of the SARS-CoV-2 genomic 5’UTR annotated to show the ORF1 viral initiation codon AUG_266_, as well as CUG_59_ and AUG_107_ codons with their corresponding short upstream ORFs uORF1a and uORF1b. Mutations (red) introduced into SL1, SL2 and SL3 are shown in boxes. (B, C) Factor requirements for 48S complex formation on genomic (B) and subgenomic (C) SARS-CoV-2 mRNAs assayed by toe-printing. The positions of 48S complexes formed on different codons are indicated on the right. (D) The influence of destabilization of SL4 on 48S complex formation on CUG_59_. Upper panel- nucleotide substitutions (red) shown on secondary structure models of *wt* and destablized SL4. Lower panel - 48S complex formation assayed by toe-printing. The position of CUG_59_ is indicated on the left. (B-D) Lanes C, T, A, and G show the corresponding sequences derived using the same primers as for toe-printing. (E) The effect of mutations introduced into the SARS-CoV-2 5’UTR (some shown in panel A) on its activity, assayed by translation in RRL in the presence of ^35^S-Methionine. (F) The influence of RRL supplementation with eIF1 and eIF1A on initiation at CUG_59_ and AUG_107_ of SARS-CoV-2 mRNA lacking stop codons UAA_134-6_ and UAA_137-9_. (E-F) Initiation codons corresponding to translation products are indicated on the sides.

48S complexes on genomic SARS-CoV-2 RNA were reconstituted *in vitro* from individual purified translation components (40S subunits, initiation factors and Met-tRNA_i_^Met^). 48S complex formation was assayed by toe-printing which involves extension by reverse transcriptase of a primer annealed to the ribosome-bound mRNA: cDNA synthesis is arrested by the leading edge of the 40S subunit yielding toe-prints +15-17 nt downstream from the +1 nt of the P site codon. Efficient 48S complex formation on AUG_266_ was observed only in the presence of all eIFs (2, 3, 4A, 4B, 4G, 1 and 1A) and DHX29, a DExH protein that binds directly to 40S subunits and promotes initiation on structured mRNAs (Pisareva et al., 2008; Yu et al., 2011; Hashem et al., 2013) (Figure 5B, lane 10). In the absence of DHX29, 48S complexes assembled efficiently on CUG_59_ and AUG_107_ (Figure 5B, lane 9). At the same time, efficient 48S complex formation on AUG_107_ required the simultaneous presence of eIF1 and eIF1A, whereas 48S complex on CUG_59_ formed efficiently with eIF1 in the absence of eIF1A (Figure 5B, lane 7). In contrast to genomic RNA, 48S complex formation on SARS-CoV-2 sgRNAs did not require DHX29, and no 48S complexes formed on CUG_59_ in this case (Figure 5C). To investigate whether the downstream SL4 (which is lacking in sg RNAs) stabilizes 48S complex formation on the near-cognate CUG_59_ of genomic RNA, we investigated assembly of 48S complexes on mutant genomic RNAs with destabilized SL4. Partial (at the base) or complete destabilization of SL4 (Figure 5D, upper panel) impaired 48S complex formation on CUG_59_ (Figure 5D, lower panel).

Next, we investigated translation in rabbit reticulocyte lysate (RRL) of mRNAs containing wt and mutant SARS-CoV-2 genomic 5’UTRs with a short adjacent coding region linked to a NanoLuc reporter. Stabilization but not disruption of SL1 (Figure 5A) strongly impaired translation of SARS-CoV-2 mRNA (Figure 5E, left panel). Stabilization of SL2 and SL3 (Figure 5A) had only a small effect on initiation at AUG_266_ (Figure 5E, right panel, lane 7). Elimination of CUG_59_ and AUG_107_ codons individually or in combination did not reduce viral translation (Figure 5E, right panel, compare lanes 1 with lanes 2, 5 and 9), indicating that reinitiation after their short ORFs does not make a meaningful contribution to initiation at AUG_266_. Mutation of stop codons of short ORFs for CUG_59_ and AUG_107_ resulted in translation of longer products, indicated that these codons can also function in RRL (Figure 5E, right panel, lanes 3 and 4). Translation from CUG_59_ in RRL seemed to be relatively less efficient than 48S complex formation on this codon during *in vitro* reconstitution. This could be due to discrimination against CUG_59_ during later stages of ribosomal subunit joining or elongation. As expected, supplementation of RRL with eIF1A, and particular with eIF1, strongly reduced initiation at CUG_59_ and AUG_107_ (Figure 5F).

Taken together, these data indicate that initiation on SARS-CoV-2 genomic RNA occurs by 5’end dependent leaky scanning which also requires DHX29.

### Properties of the SARS-CoV-2 5’UTR accounting for its resistance to Nsp1-mediated cleavage

Resistance of SARS-CoV-2 to Nsp1-mediated inhibition was attributed to the presence of the 5’-terminal SL1 (Banerjee et al., 2020; Mendez et al., 2021; Tidu et al., 2021; Vora et al., 2022). We therefore compared Nsp1-mediated cleavage of wt and SL1 mutant mRNAs (Mutants 1-3; Figure 6A, upper panel). To preserve the overall structure of SL1, we swapped 5’ and 3’ strands in the SL1A and SL1B stems individually or in combination. To induce cleavage, wt and mutant mRNAs were incubated with Nsp1, 40S subunits and eIFs, after which cleavage sites were identified by primer extension. SL1 contains two pairs of guanosines: G_7_G_8_ and G_23_G_24_. The pair is too close to the 5’end to induce cleavage, whereas the other pair, on the other hand, were suggested to be too far away. Consistently, no cleavage was observed in the case of *wt* mRNA (Figure 6A, lower panel, lane 2). Swapping strands in SL1A in Mutant 1 led to the appearance of Gs at positions 32-33, which resulted in the appearance of the unexpected weak cleavage between A_4_A_5_ (Figure 6A, lower panel, lane 4). Swapping strands in SL1B in Mutant 2 brought two Gs to positions 15-16, which promoted regular cleavage at upstream nucleotides (Figure 6A, lower panel, lane 6). Swapping strands in SLA1 and SL1B simultaneously in Mutant 3 also induced canonical cleavage upstream of G_15_G_16_ and G_32_G_33_ (Figure 6A, lower panel, lane 8). Since cleavage directed by G_32_G_33_ did not occur in Mutant 1, its appearance in Mutant 3 almost certainly resulted from prior cleavage directed by G_15_G_16_, which brought G_32_G_33_ closer to the 5’ end (positions 23-24) in the truncated RNA. Weak cleavage between A_4_A_5_, which was observed Mutant 1, also occurred in Mutant 3. A possible explanation for this cleavage is the accommodation in the active center of only partially unwound SL1. Taken together, these data indicate that cleavage is not influenced by SL1 secondary structure because bringing Gs close to the 5’end in Mutants 2 and 3 without destabilization of SL1 induced Nsp1-mediated cleavage. However, it raises the question of why G_23_G_24_ in the wt SL1 do not induce cleavage, whereas GG at the same positions in Mutant 3 do.

**Figure 6.**
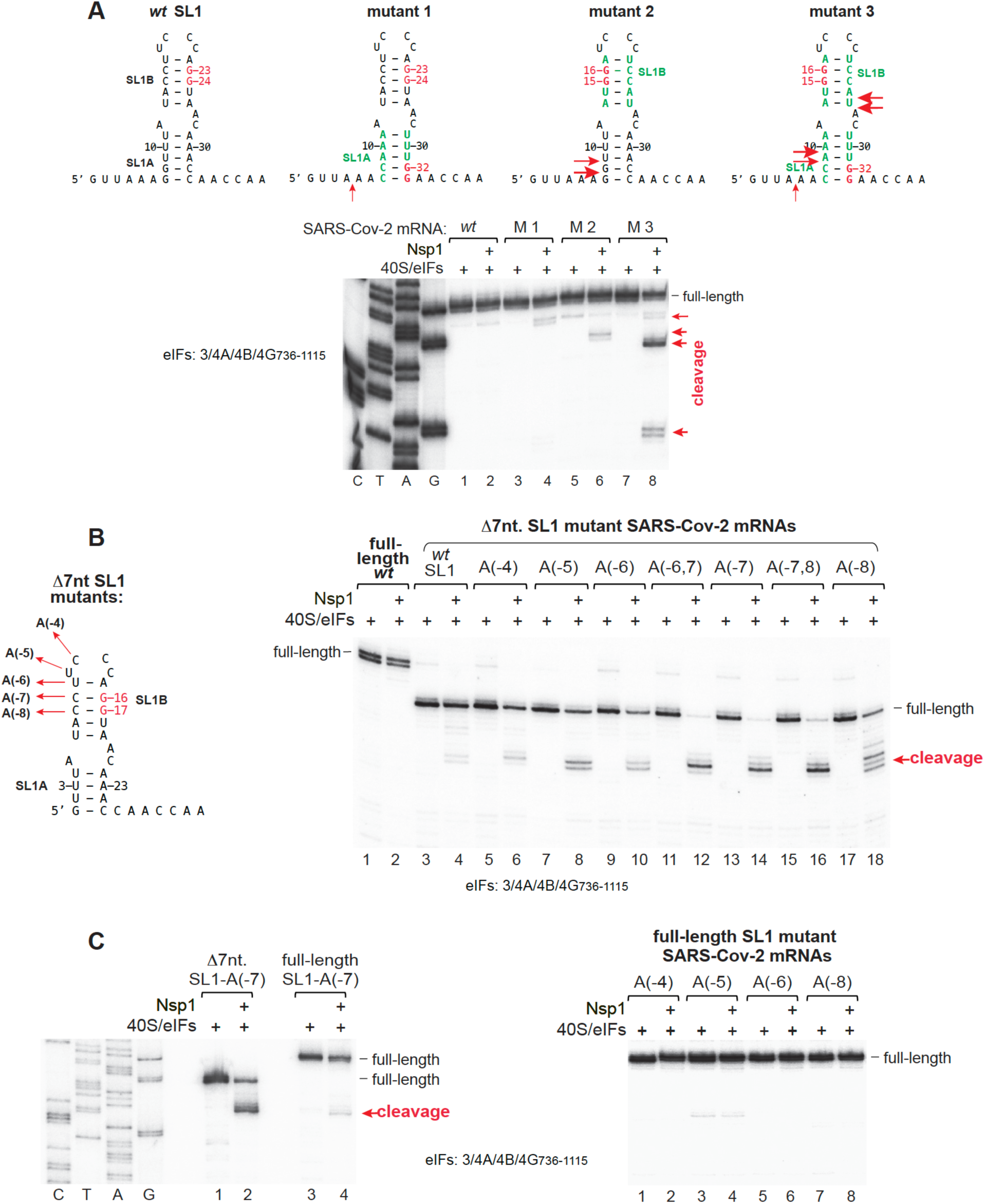
Nsp1-mediated cleavage of SARS-CoV-2 mRNAs. (A) Upper panel - secondary structure models of the *wt* and mutant SARS-CoV-2 SL1. G nucleotides are in red, introduced mutations are in green. Lower panel - Nsp1-mediated cleavage of *wt* and mutant SARS-CoV-2 mRNAs shown in the upper panel in the presence of the indicated translation components, assayed by primer extension. Cleavage sites are indicated on the right and marked on the SL1 models shown in the upper panel (red arrows). Lanes C, T, A, and G show the *wt* sequence derived using the same primer as for primer extension. (B) Left panel – mutations introduced into SL1 of the 5’-terminally truncated (D7 nt.) SARS-CoV-2 5’UTR. Right panel - Nsp1-mediated cleavage of *wt* and mutant SARS-CoV-2 mRNAs (shown in the left panel) in the presence of the indicated translation components, assayed by primer extension. Cleavage sites are indicated on the right (red arrows). (C) Nsp1-mediated cleavage of full-length SARS-CoV-2 mRNAs containing mutations identical to those that were introduced into the 5’-terminally truncated SARS-CoV-2 5’UTR (B, left panel). mRNAs were incubated with Nsp1 and the indicated translation components and assayed by primer extension. The cleavage site is indicated on the right. Lanes C, T, A, and G show the *wt* sequence derived using the same primer as for primer extension.

We noticed that in addition to being relatively far from the 5’ end, G_23_G_24_ in SL1 are preceded by a long oligoPy stretch. Therefore, to investigate the relative influence of the position and the nature of the preceding nucleotide region on Nsp1-mediated cleavage of SARS-CoV-2 mRNA, we 5’-terminally truncated this RNA by 7 nucleotides and introduced Py to A substitutions at unique positions upstream of G_23_G_24_ (Figure 6B, left panel). Truncation of SARS-CoV-2 mRNA by 7 nucleotides resulted in susceptibility to Nsp1, but cleavage was very weak (Figure 6B, right panel, lane 4). Introduction of a single A at positions -4, -5 or -6 moderately increased cleavage efficiency (Figure 6B, lanes 6, 8 and 10), whereas introduction of A at position -7 resulted in complete cleavage of RNA (Figure 6B, lane 14). With A at position -8, cleavage occurred at several sites (Figure 6B, lane 18). Introduction of the same Py to A substitutions into full-length SARS-CoV-2 mRNA resulted in cleavage of RNA containing A at position - 7 (Figure 6C, left panel), but not of mRNAs with A at other positions (Figure 6C, right panel). These data again underscore the importance of the Pu nucleotide at position -7 upstream of the critical guanosine.

Taken together, our data indicate that the resistance to Nsp1-mediated cleavage of SARS-CoV-2 mRNA is caused by both the distance of G nucleotides from the 5’end and the nature of the upstream nucleotides.

### The influence of Nsp1 on translation initiation on the SARS-CoV-2 5’UTR and its ability to compete with cellular mRNAs

Although SARS-CoV-2 mRNA is not susceptible to Nsp1-mediated cleavage, like most cellular mRNAs, it is translated following initiation by the 5’end dependent scanning mechanism and should therefore also be inhibited by binding of the C-terminal domain of Nsp1 into the entrance portion of the mRNA-binding channel where it interferes with binding of mRNA (Schubert et al., 2020; Thoms et al., 2020; Yuan et al., 2020). Consistently, Nsp1 inhibited 48S complex assembly on genomic and ORF8 subgenomic SARS-CoV-2 mRNAs (Figure 7A) and inhibited translation in RRL of mRNA containing SARS-CoV-2 genomic 5’UTR in a dose-dependent manner (Figure 7B).

**Figure 7.**
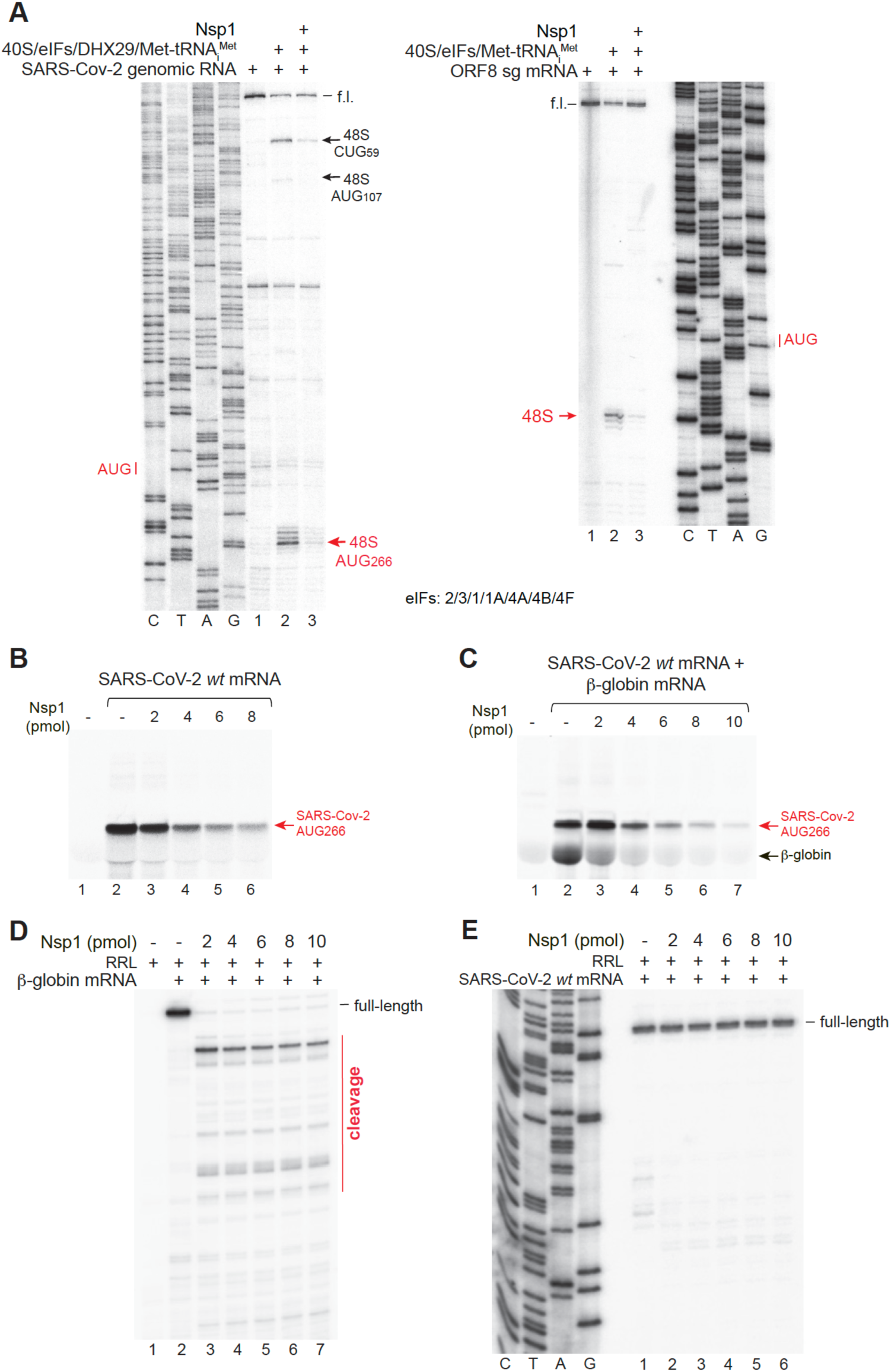
The influence of Nsp1 on initiation on SARS-CoV-2 mRNA. (A) The influence of Nsp1 on 48S complex formation on mRNAs containing genomic and ORF8 subgenomic SARS-CoV-2 5’UTRs in the presence of the indicated translation components, assayed by toe-printing. The positions of 48S complexes are indicated. (B) The influence of Nsp1 on translation of mRNA containing the genomic SARS-CoV-2 5’UTR in RRL supplemented with the indicated amounts of Nsp1, assayed in the presence of ^35^S-Methionine. (C) The influence of Nsp1 on competition between b-globin mRNA and mRNA containing the genomic SARS-CoV-2 5’UTR during translation in RRL supplemented with the indicated amounts of Nsp1, assayed in the presence of ^35^S-Methionine. (B, C) Translation products resolved by SDS-PAGE are indicated on the right. (D, E) The integrity of (D) b-globin mRNA and (E) mRNA containing the genomic SARS-CoV-2 5’UTR after translation of their mixture in RRL supplemented by the indicated amounts of Nsp1. After translation, mRNAs were phenol extracted, ethanol precipitated and assayed by primer extension. Cleavage sites (in D) are shown on the right (red).

This poses the question of how SARS-CoV-2 genomic and subgenomic mRNAs outcompete cellular mRNAs in cells expressing Nsp1. For this, we investigated competition between mRNA containing the SARS-CoV-2 genomic 5’UTR and native β-globin mRNA during translation in RRL in conditions of increasing concentrations of Nsp1. Although at higher concentrations Nsp1 inhibited translation of both mRNAs, at a lower concentration, it inhibited translation of only β-globin mRNA, whereas translation of mRNA containing SARS-CoV-2 genomic 5’UTR increased (Figure 7C). Analysis of mRNA integrity revealed that even at the lowest concentration of Nsp1, β-globin mRNA was completely cleaved (Figure 7D), whereas SARS-CoV-2 mRNA remained intact at all Nsp1 concentrations (Figure 7E). These data suggest that during infection, those ribosomal preinitiation complexes which are associated with Nsp1 would induce rapid cleavage of cellular mRNAs, allowing SARS-CoV-2 mRNA to be very efficiently translated by preinitiation complexes lacking Nsp1 without any competition with cellular mRNAs.

## DISCUSSION

Using the *in vitro* reconstitution approach, we determined that SARS-CoV-2 Nsp1-mediated endonucleolytic cleavage of RNA, which strictly requires 40S subunits and eIF3g, proceeds via a transesterification mechanism which involves intramolecular nucleophilic attack of the 2’OH of the ribose on the adjacent phosphodiester bond to yield 5’OH and 2’,3‘-cyclic phosphate-linked cleavage products. Cleavage absolutely depends on the presence of a guanosine residue within the first ∼10-25 5’-terminal mRNA nucleotides and occurs in a specific window upstream of this G, with cleavage between -6/-7 and - 7/-8 nucleotides relative to the G being most efficient. It also shows cleavage site preference for Pu (preferably followed by Py) at positions -7 or -8 whose 2’OH hydroxyl group would be positioned for the nucleophilic attack on the adjacent phosphodiester bond. In the absence of Pu (or PuPy pairs) at the optimal distance from G, lower-efficiency cleavage occurs at multiple sites, either at the optimal distance (but between other nucleotide pairs), or between PuPy pairs located at a suboptimal distance from the G. Cleavage was least efficient when the G was preceded by extended oligoPy stretches. The cleavage closest to and most distant from the G occurred between -5/-6 positions and -9/-10 nucleotides, respectively, establishing a 4-5 nt-long window for cleavage upstream of the critical G. Thus, the presence of a guanosine near the 5’ end of RNA is an absolute prerequisite for Nsp1-mediated cleavage, whereas requirements for the specific nucleotides at the cleavage site and for its precise position are more relaxed.

The absolute dependence of Nsp1-mediated cleavage on the presence of a guanosine near the 5’ end of RNA explains previous observations that a guanosine-free window close to the 5’ end confers resistance to Nsp1-mediated translational shut off for both cellular and viral mRNAs (Slobodin et al., 2022; Chen et al., 2023; Berlanga et al., 2025; Galbraith et al., 2026). Importantly, given the relatively large region downstream from the 5’ end in which the presence of a guanosine can sensitize mRNA to Nsp1-mediated cleavage and the relatively relaxed requirements for the specific nature and position of the cleavage site relative to the critical G residue, the vast majority of cellular mRNAs would be susceptible to Nsp1-mediated cleavage and translational shut off.

Resistance of SARS-CoV-2 mRNA to Nsp1-mediated translational inhibition was shown to depend on the absence of G residues in a window 9-22 nt from its 5’ end (Slobodin et al., 2022; Chen et al., 2023; Berlanga et al., 2025). However, in addition to being relatively far from the 5’ end, G_23_G_24_ in SL1 of SARS-CoV-2 are preceded by a long oligoPy stretch with two pyrimidines (CC) located at the important positions -7 and -8 from G_23_G_24_. We found that bringing these Gs closer to the 5’ end by truncation of SARS-CoV-2 mRNA by 7 nucleotides or substituting C by A at position -7 relative to G_23_ in the full-length mRNA both led to Nsp1-induced cleavage of moderate intensity, whereas simultaneous introduction of both changes resulted in very efficient cleavage. Thus, resistance to Nsp1-induced cleavage of SARS-CoV-2 mRNA is ensured at two levels: the primary requirement for recognition of the essential guanosine and the influence of the nucleotide context of the cleavage site. In SL1 of SARS-CoV mRNA, which like SARS-CoV-2 mRNA is resistant to Nsp1-mediated cleavage (Figure S2), G_23_G_24_ are also preceded by an oligoPy stretch with a CC dinucleotide at positions -7 and -8. In contrast, the key resistance features are lacking in MERS-CoV SL1, and this mRNA underwent efficient Nsp1-induced cleavage (Figure S2). Consistently, although MERS-CoV Nsp1 binds to the 40S subunit and inhibits translation (Devarkar et al., 2023; Maurina et al., 2023), it does so without triggering RNA degradation (Bäumlin et al., 2025).

Using an *in vitro* reconstitution approach, we confirmed that initiation on SARS-CoV-2 mRNA occurs by 5’ end-dependent leaky scanning past AUG_107_ and the near-cognate CUG_59,_ which is in line with the strong dependence of SARS-CoV-2 translation on the eIF4F cap-binding complex (Condé et al., 2022) and suppression of initiation at the near-cognate CUG_59_ by eIF1 and eIF1A (Aviner et al, 2024). We also found that initiation on genomic, but not on subgenomic, SARS-CoV-2 mRNA additionally requires DHX29, likely due to the presence of SL4 and SL5 in the former. Although SARS-CoV-2 mRNA was not cleaved by Nsp1, consistent with the mechanism of initiation on this mRNA, like cellular mRNAs, it was strongly inhibited by Nsp1 due to binding of its C-terminal domain to the entrance portion of the ribosomal mRNA-binding channel (Schubert et al., 2020; Thoms et al., 2020; Yuan et al., 2020). Our results concerning translational competition between SARS-CoV-2 and β-globin mRNAs in conditions of increasing concentrations of Nsp1 suggest the following mechanism for Nsp1-mediated enhancement of SARS-CoV-2 translation. Following over-expression of Nsp1 (Bujanic et al., 2022; Slobodin et al., 2022; Maurina et al., 2023) or during viral infection, 40S-associated Nsp1 would induce rapid cleavage of cellular mRNAs and consequent abrogation of their translation, whereas SARS-CoV-2 mRNA would be efficiently translated without competition from cellular mRNAs by Nsp1-free 43S preinitiation complexes.

The strict requirements of 40S subunits and eIF3g for Nsp1-mediated RNA cleavage raises the question of the exact roles of all three components in recognition of the critical guanosine and in activation of the 2’OH in the nucleotide at positions -7 or -8 for the nucleophilic attack on its 3’ phosphate group. Our attempt to employ specific zero-length UV crosslinking to determine the component which directly interacts with the critical guanosine was not successful because replacement of G by 6thioG abrogated Nsp1-mediated cleavage, indicating that this modification impairs specific recognition of guanosine by e.g. disrupting the pattern of hydrogen bonds or by steric clashes induced by the larger sulfur atom. However, UV crosslinking of RNAs with 4thioU at different positions relative to the G revealed efficient specific 40S subunit-dependent interactions of Nsp1 with nucleotides at positions -1 and -2 which were stimulated by eIF3g to different extents. Thus, whereas efficient crosslinking of Nsp1 to position -1 occurred even without eIF3g, crosslinking to position -2 was completely eIF3g-dependent. Weak eIF3g-dependent crosslinking of Nsp1 was also observed for positions -3 and -4. Exclusive crosslinking of Nsp1 to -1 and -2 nucleotides that are directly neighboring to the critical G suggests that Nsp1 might be primarily responsible for recognition of the guanosine. KR72-73AA and E91A substitutions abrogated crosslinking of Nsp1 to both -1 and -2 positions. Given the different dependence of crosslinking at these positions on eIF3g, these results suggest that these amino acids could be involved in specific recognition of guanosine and/or the immediately preceding nucleotides rather than in a putative Nsp1-eIF3g interaction. The absolute requirement for eIF3g in paradigm for the virus-induced suppression of cellular translation during infection cross-linking of Nsp1 to position -2, -3 and -4 suggests that eIF3g’s essential role could be to promote routing of the upstream nucleotides into the cleavage center. Alanine substitutions of some Nsp1 residues that were essential for Nsp1 cleavage (e.g. RK124-125, K129 or H13) (Abaeva et al., 2023) reduced but did not abrogate Nsp1 crosslinking to positions -1 and -2, indicating that these residues could be involved in binding to nucleotides that are further upstream or in activation of 2’OH. Unfortunately, UV cross-linking did not reveal any specific contacts of positions -9, -8, -6 and -5 that surround the actual cleavage site.

Nsp1-mediated cleavage is not the only example of an endoribonucleolytic process that is coupled to translation. Other ribosome-dependent endonucleases that act using the transesterification mechanism include, for example, RelE which cleaves mRNA after the second nucleotide in the ribosomal A site (Pedersen et al., 2003; Neubauer et al., 2009) and ANKZF1 which induces specific cleavage in the tRNA acceptor arm, acting on 60S ribosome-nascent chain complexes to release proteasome-degradable ubiquitinated nascent chains linked to three 3’-terminal tRNA nucleotides (Kuroha et al., 2018; Su et al., 2019; Yip et al., 2019). In both cases, ribosomes interact directly with RNA substrates in the area of cleavage. By contrast, we did not detect any specific UV crosslinking of RNA with 40S subunits in Nsp1-associated cleavage complexes. Moreover, the 40S-RNA crosslinking that presumably occurred by binding of unstructured RNA to the mRNA-binding channel was strongly inhibited by Nsp1 which is consistent with binding of Nsp1 to the mRNA-binding channel and blocking its ability to associate with the RNA. The role of the 40S subunit during Nsp1-mediated cleavage could therefore be both to serve as a scaffold that brings Nsp1 and eIF3g together and to influence the conformation of Nsp1. Future identification of the exact roles of Nsp1 and eIF3g (and their specific residues) in activation of 2’OH as well stabilization of the transition state and 5’OH leaving group will require determination of the structure of the 40S/Nsp1/eIF3g/RNA complex.

## MATERIALS AND METHODS

### Construction of plasmids

Vectors for bacterial expression of eIF1, eIF1A, eIF3g, eIF4A, eIF4B, eIF4G_736-1115,_ *E. coli* methionyl tRNA synthetase, DHX29 and *wt* and mutant forms of His_6_-Nsp1(SARS-CoV-2), and for transcription of tRNA_i_^Met^ have been described (Abaeva et al., 2023; Pestova et al., 1996a,b; Pestova et al., 1998; Pestova and Hellen, 2001; Lomakin et al., 2006; Dhote et al., 2012).

Transcription vectors for mRNAs containing 5’UTRs comprising CAA repeats with various mutations (Figures 1A, 3 and 4A-B) were based on the MSHL-STOP vector (Zinoviev et al., 2018) and consisted of DNA for a T7 promoter followed by a (CAA)n-derivatised 5’UTR, an MSHL short ORF, a stop codon and nt 16-121 of the rabbit β-globin ORF cloned between EcoRV and HindIII sites of pUC57 (GenScript, Piscataway, NJ, USA).

SARS-CoV-2(nt.2-301)-nLUC-3UTR_pUC57 for transcription of mRNAs employed for *in vitro* reconstitution of 48S initiation complexes and Nsp1-mediated cleavage was made by GenScript by inserting DNA between EcoRV and EcoRI sites of pUC57 that corresponded to a T7 promoter followed by a single G residue, SARS-CoV-2 nt 2-301 and nt 29671-29867 (GenBank: NC_045512) flanking an in-frame nanoluciferase (nLuc) open reading frame (GenBank: KM359774.1). This vector was used to generate constructs with mutations in SL1 and SL4 (Figures 5D, 6A).

SARS-CoV-2(nt.2-361)-nLUC-3UTR_pUC57, which was used to transcribe mRNAs for *in vitro* translation, was made by Synbio (Monmouth Junction, NJ, USA) and contained SARS-CoV-2 nt 2-361 and nt 29671-29867 (GenBank: NC_045512) flanking the nLuc ORF (GenBank: KM359774.1) that had been modified by substitutions to introduce AUG codons at triplets 20, 38, 58, 133, 144, 165 and 170 (to increase labeling during in vitro translation). It was used to generate variants described in Figures 5A and 5E-F.

The SARS-CoV transcription vector CoV(nt.2-360)-nLUC-3UTR was made by Synbio by inserting DNA between EcoRV and EcoRI sites of pUC57 that corresponds to a T7 promoter followed by a single G residue, SARS-CoV nt 2-360 (GenBank: AY278741.1), an in-frame nLuc ORF modified as described above, and SARS-CoV nt 29385-29727.

The MERS-CoV transcription vector MERS-Cov(nt.2-374)-nLUC-3UTR was made by Synbio by inserting DNA between EcoRV and EcoRI sites of pUC57 that corresponded to a T7 promotor followed by a single G residue, MERS-CoV nt. 1-374 (NCBI Ref. seq. NC_019843.3), an in-frame nLuc ORF modified as described above, and MERS-CoV nt. 29805- 30108.

Vectors for transcription of SARS-CoV-2 sg mRNAs were made by GenScript and were based on sg mRNA sequences (Kim et al., 2021; Wang et al., 2021). The Spike sg mRNA pUC57-based vector consisted of EcoRV and HindIII restriction sites flanking a T7 promoter, a G nucleotide and SARS-CoV-2 nt 2-65 linked to nt 21551-21863. The analogous Membrane and ORF8 sg mRNA transcription vectors contained a T7 promoter, a G nucleotide and SARS-CoV-2 nt 1-64 linked to 26467-26823 or SARS-CoV-2 nt 1-65 linked to nt 27883-28304, respectively, cloned between EcoRV and HindIII restriction sites in pUC57. The pUC57-sgORF8_wt vector was used to generate derivatives described in Figures 6B-C.

Vectors for transcription of CrPV nt 5997-6320 (NCBI Ref. seq. NC_003924.1) corresponding to the IRES and the adjacent cognate coding sequence or mutant variants thereof with substitutions at positions flanking the primary cleavage sites at nt 6246-7 (Figure 4C) were made by GenScript. DNA comprising a T7 promoter, *wt* or mutant viral sequences, an upstream BamHI restriction site and downstream EcoRV and EcoRI restriction sites was cloned between BamHI and EcoRI sites of pUC57.

### Preparation of RNAs

All mRNAs were *in vitro* transcribed using T7 RNA polymerase (ThermoFisher). Custom 14-30 nt.-long RNA oligonucleotides and their derivatives (Figures 1B-F, 2, S1) were made by Horizon Discovery Biosciences (Lafayette, CO, USA). They were 5’[^32^P]-labeled by T4 polynucleotide kinase (PNK) (New England BioLabs, Ipswich, MA, USA) using [γ-^32^P]ATP (222 TBq/mmol) (Revvity, Waltham, MA, USA). After phosphorylation, T4 PNK was inactivated by incubating reaction mixtures at 65°C for 20 minutes.

### Purification of ribosomal subunits, initiation factors, *E. coli* methionyl tRNA synthetase and recombinant Nsp1

Native mammalian 40S ribosomal subunits, eIF2, eIF3 and eIF4F were purified from rabbit reticulocyte lysate (RRL) (Green Hectares, Oregon, WI) as described (Pisarev et al., 2007). Human recombinant eIF1, eIF1A, eIF4A, eIF4B, eIF4G_736-1115_, eIF3g and DHX29, *E. coli* methionyl-tRNA synthetase, and *wt* and mutant SARS-CoV-2 Nsp1 were expressed in *E. coli* BL21(DE3) (Invitrogen, Carlsbad, CA) and purified as described (Pestova et al., 1996a,b; Pestova et al., 1998; Lomakin et al., 2006; Dhote et al., 2012; Abaeva et al., 2023). *In vitro* transcribed tRNA_i_^Met^ was aminoacylated using recombinant *E. coli* methionyl-tRNA synthetase (Lomakin et al., 2006; Pisarev et al., 2007).

### *In vitro* reconstitution of 48S complexes on SARS-CoV-2 mRNAs

For assembly of 48S initiation complexes on genomic and subgenomic SARS-CoV-2 mRNAs, 1 pmol RNA was incubated with 2 pmol 40S ribosomal subunits, 3 pmol Met-tRNA_i_^Met^ and indicated combinations of 6 pmol eIF2, 4 pmol eIF3, 8 pmol eIF4A, 2 pmol eIF4B, 2 pmol eIF4F, 10 pmol eIF1, 10 pmol eIF1A and 0.3 pmol DHX29 for 10 min at 37°C in 40 μl of buffer A containing 20 mM Tris-HCl (pH 7.5), 100 mM KCl, 2.5 mM MgCl₂, 2 mM DTT, 0.25 mM spermidine, 1 mM ATP, 0.4 mM GTP and 1.5 U/μl RiboLock RNAse inhibitor. To assay the influence of Nsp1 on 48S complex formation, 40S subunits were preincubated with 10 pmol Nsp1 for 5 min at 37°C before addition of other components. 48S complex assembly was assayed by toe-printing using AMV reverse transcriptase (RT) (Promega) and ^32^P-labeled primer (Pisarev et al., 2007). Radiolabeled cDNAs were phenol-extracted, ethanol-precipitated, resolved on 6% polyacrylamide sequencing gel and visualized by phosphoimager.

### Nsp1-induced cleavages on long mRNAs

To reconstitute Nsp1-induced cleavage of mRNA, 2 pmol 40S subunits were first preincubated with 10 pmol Nsp1 for 5 min at 37°C in 40 μl of buffer A, then reaction mixtures were supplemented with 1 pmol RNA, 4 pmol eIF3, 8 pmol eIF4A, 2 pmol eIF4B and 6 pmol eIF4G_736-1115_ (or just with 5 pmol eIF3g in the case of the CrPV IRES) after which incubation continued for 10 more minutes. To detect cleavage, mRNAs were analyzed by primer extension using AMV RT and ^32^P-labeled primer (Pisarev et al., 2007). Radiolabeled cDNAs were phenol-extracted, ethanol-precipitated, resolved on 6% polyacrylamide sequencing gel and visualized by phosphoimager.

### Analysis of Nsp1 cleavage on short RNAs

1 pmol of 5’[^32^P]-labeled RNA was incubated with 2 pmol 40S ribosomal subunits, 10 pmol Nsp1 and different combinations of 10 pmol eIF3g, 5 pmol eIF3, 2 pmol eIF4B, 8 pmol eIF4A, and 6 pmol eIF4G_736-1115_ in 10 μl of buffer A for 5 min at 37°C. Where indicated, 10 U of T4 PNK was added after incubation, and the reaction mixture was incubated for an additional 10 min. RNA cleavage products were directly resolved on a 20% polyacrylamide sequencing gel and visualized by phosphorimager.

### UV-crosslinking

Different combinations of 4 pmol 40S ribosomal subunits, 5 pmol Nsp1 and 5 pmol eIF3g were preincubated for 5 min at 37°C in 20 μl buffer A. After addition of 1 pmol 5’[^32^P]-labeled thioU-containing RNA, reactions were incubated for 2 min at room temperature, placed on ice, and UV-irradiated at 360 nm for 25 min using a Stratagene UV Stratalinker 1800. Crosslinked complexes were then directly resolved on NuPAGE 4–12% Bis-Tris precast gels (Invitrogen) and visualized by phosphoimager.

### *In vitro* translation in rabbit reticulocyte lysate (RRL)

1 pmol of SARS-CoV-2 mRNA individually or in combination with 2 pmol native β-globin mRNA was translated in RRL (Promega) at 30°C for 30 min in 20 μl reaction mixtures containing [^35^S]methionine (Revvity) and the indicated amounts of Nsp1. Translation products were analyzed by electrophoresis on NuPAGE 4–12% Bis-Tris precast gels. The integrity of phenol-extracted and ethanol-precipitated mRNAs was by assayed by primer extension using AMV RT and ^32^P-labeled primer (Pisarev et al., 2007). Radiolabeled cDNAs were phenol-extracted, ethanol-precipitated, resolved on 6% polyacrylamide sequencing gel and visualized by phosphoimager.

## ACKNOWLEDGMENTS

This work was supported by NIH grant GM122602 (to T.V.P), and NIH grant AI166944 to (C.U.T.H and T.V.P).

## AUTHOR CONTRIBUTIONS

I.S.A. and A.B.J. performed the experiments. T.V.P., C.U.T.H. and I.S.A. designed experiments and interpreted data. T.V.P. and C.U.T.H. wrote the paper with the input from all authors.

